# An IPI based immune prognostic model for diffuse large B-cell lymphoma

**DOI:** 10.1101/2021.03.03.433839

**Authors:** Shidai Mu, Deyao Shi, Lisha Ai, Fengjuan Fan, Fei Peng, Chunyan Sun, Yu Hu

## Abstract

**Background:** International Prognostic Index (IPI) was widely used to better discriminate prognosis of patients with diffuse large B-cell lymphoma (DLBCL). However, there is a significant need to identify novel valuable biomarkers in the context of targeted therapies, such as immune checkpoint blockade (ICB) therapy.

**Methods:** Gene expression data and clinical information of DLBCL were obtained from The Cancer Genome Atlas (TCGA) and Gene Expression Omnibus (GEO) datasets. 371 immune-related hub genes in DLBCL patients with different IPI levels were identified by weighted gene co-expression network analysis (WGCNA), and 8 genes were selected to construct an IPI-based immune prognostic model (IPI-IPM). Afterward, the genetic, somatic mutational and molecular profiles of IPI-IPM subgroups were analyzed, as well as the potential clinical response of ICB in different IPI-IPM subgroups.

**Results:** The IPI-IPM was constructed based on the expression of CMBL, TLCD3B, SYNDIG1, ESM1, EPHA3, HUNK, PTX3 and IL12A, where high-risk patients had shorter overall survival (OS) than low-risk patients, consistent with the results in the GEO cohorts. The comprehensive results showed that high IPI-IPM risk scores were correlated with immune-related signaling pathways, high KMT2D and CD79B mutation rates, as well as up-regulation of inhibitory immune checkpoints including PD-L1, BTLA and SIGLEC7, indicating more potential response to ICB therapy.

**Conclusion:** The IPI-IPM has independent prognostic significance for DLBCL patients, which provides an immunological perspective to elucidate the mechanisms on tumor progression, also sheds a light on developing immunotherapy for DLBCL.

## Introduction

Diffuse large B-cell lymphoma (DLBCL) accounts for about 40% of non-Hodgkin B-cell lymphoma (NHL), with an annual incidence rate of over 100,000 cases worldwide^1, 2^. Although current frontline DLBCL therapy (the standard R-CHOP chemotherapy regimen) is associated with a high complete response rates of 70–80%, 10% to 15% of DLBCL patients are refractory, and almost 40% of cases experience relapse within 2–3 years after initial response^3, 4^. With the development of high-throughput technologies, germinal center B-cell-like (GCB) and activated B-cell-like (ABC) DLBCL subtypes were identified by gene expression profiling (GEP) based on cell-of-origin (COO) classification^5-7^. More recently, several key cytogenetic alterations including mutations, somatic copy number alterations (SCNA) and structural variants (SV), have been shown to classify distinct genetic subtypes within the COO subgroups, providing insights into disease heterogenous pathogenesis and candidate treatment targets^1, 3, 7-9^. Several prognostic factors including COO and the International Prognostic Index (IPI) have already been identified in the rituximab era, which still need to be further investigated in the context of targeted therapies^7, 9-12^. Therefore, there is an urgent need to explore potential molecular mechanism and identify more key biomarkers and therapeutic targets.

Accumulating evidence has shed light on the prognostic role of tumor microenvironments (TME) in immune checkpoint blockade therapy (ICB), which was mostly composed of a variety of immune cells (T-, NK-, and B-cells as well as macrophages) and stroma (blood vessels and extra-cellular matrix)^13-16^. Kotlov et al. characterized the DLBCL TME into 4 distinct microenvironment compositions including “germinal center-like” (GC), “mesenchymal” (MS), “inflammatory” (IN) and “depleted” (DP) form, which associate with distinct clinical behavior and provide novel potential targets for innovative therapeutic interventions^11^. In this study, we identified immune-related hub genes in DLBCL patients at different IPI levels by weighted gene co-expression network analysis (WGCNA), and constructed an IPI-based immune prognostic model (IPI-IPM). We then characterized the genetic, somatic mutational and molecular profile of IPI-IPM subgroups, investigated the expression of several inhibitory immune checkpoints between low- and high-risk subgroups, and applied. The results showed that IPI-IPM was a promising prognostic biomarker, which also had potential for use in patient management.

## Materials and Methods

### Data selection and acquisition

Data acquisition of the present study is fully under the TCGA publication guidelines (https://www.cancer.gov/tcga). Gene expression data (RNA-seq) of 570 samples, masked somatic mutation data of 37 samples and clinical follow up data with clinicopathological characteristics of 566 patients of DLBCL projects (TCGA-DLBC, CTSP-DLBCL1, NCICCR-DLBCL) were obtained from the Cancer Genome Atlas (TCGA) by using the TCGAbiolinks^17^ R package in R software (version 4.0.2, https://www.r-project.org). Clinical information, gene expression subtype and genetic subtype of the DLBCL patients were supplemented by referring to open-access supplementary files of the GDC DLBCL publication^4^. Matched gene expression data and survival follow-up data can be obtained in 563 samples. Matched gene expression data and survival follow-up data with available IPI data can be obtained in 445 samples (**Supplementary Table 1**). Gene expression data (Microarray) with matched clinical information of 414 DLBCL patients were obtained from GSE10846^12, 18^ by accessing the GEO database (www.ncbi.nlm.nih.gov/geo). The immunologic gene lists used in the present study were downloaded from the ImmPort (www.immport.org/shared/home) and ImmuneSigDB^19^ through MSigDB Collections (www.gsea-msigdb.org/gsea/msigdb). For the RNA-Seq data, HTSeq-count data was downloaded. Combat-Seq function of Sva^20^ R package was used to remove batch-effect among different projects. The principal components analysis was performed and visualized to examine the batch effect^21^. For each gene, the effective gene length was extracted by using EDASeq^22^ R package and a TPM (Transcripts Per Kilobase Million) gene expression data was calculated through the count2TPM function utilized in the IOBR^23^ R package by using the batch-free count data and corresponding effective gene length. Homo sapiens GRCh38 annotation file downloaded from Ensembl^24^ was used for gene symbol annotation. DESeq2^25^ R package was applied for the normalization of RNA-seq counts data and variance stabilizing transformation (VST) data was used for downstream analysis. DLBCL microenvironmental (LME) signatures and LME subtype categorizing method were referred to Kotlov et al.’s publication on Cancer Discovery^11^. An anti-PD-L1 treatment cohort (IMvigor210 cohort) with the gene expression and overall survival information were downloaded from http://research-pub.gene.com/IMvigor210CoreBiologies^26^.

### Identification of differentially expressed genes

Samples with available IPI information were categorized to high, intermediate and low risk groups according to criterion of IPI, and the DESeq2^25^ R package was applied to identify differentially expressed genes (DEGs) between high and low risk groups. The differential expression was defined with a fold-change of threshold at 1.5 and a false discovery rate (FDR) value < 0.05.

### Gene functional enrichment analysis

The clusterProfiler^27^ R package was used for both over representation analysis and pre-ranked gene set enrichment analysis (GSEA). Analysis of Gene Ontology (GO)^28^, Kyoto Encyclopedia of Genes and Genomes (KEGG) pathway^29^, and Reactome pathway^30^ was contained in the present study. Adjusted P value<0.05 was considered statistically significance. The threshold for GSEA was set at the p-value < 0.05, FDR < 0.05 and | normalized enrichment score (NES) | > 1.0. The non-parametric gene set variation analysis was further performed with the GSVA^31^ package of R.

### Weighted gene co-expression network analysis

Weighted gene co-expression network analysis (WGCNA) is commonly used for analyzing high-throughput gene expression data with different characteristics, so as to mine gene co-expression networks and intramodular hub genes based on pairwise correlations in genomic applications. In the present study, we applied the WGCNA^32, 33^ R package to analyze key gene clusters that were most relevant to IPI scores in DLBCL samples.

### Construction and validation of IPI-based immune prognostic model (IPI-IPM)

IPI-based immune related genes were selected to construct the prognostic risk model. The training cohort (563 patients with matched normalized RNA-seq data and survival data from TCGA) was used for the construction of IPI-IPM, and the testing cohort (414 patients with matched normalized microarray data and survival data from GSE10846) was used for validation of the prognostic risk model. The Survival R package was used to analyze the correlation between the expression of objective gene sets and DLBCL patients’ overall survival (OS). Univariate Cox regression analysis was performed to screen genes, of which the expression was associated with OS with a P value < 0.05. Lasso (least absolute shrinkage and selection operator) regression analysis was applied for variable selection and regularization to enhance the prediction accuracy and interpretability by using the glmnet^34^ R package. Multivariate Cox regression analysis was then carried out to select the optimal genes, based on the method of Akaike information criterion (AIC)^35^. For each sample, the risk score equals the sum of the normalized expression of each gene multiplying its corresponding regression coefficient. Time-dependent ROC curves were plotted by using the survivalROC^36^ R package. 563 DLBCL patients in the training cohort were divided to low- and high-risk score groups according to the optimal cut-off value with largest AUC in the receiver operating characteristic (ROC) curve of the median survival time. Then, Kaplan-Meier survival analysis and time-dependent ROC curve analysis were performed to evaluate the prognostic significance and accuracy of IPI-IPM^37^. Besides, Harrell’s concordance index (C-index) was calculated by using the survcomp^38^ R package. Univariate and multivariate Cox regression analyses were performed on the risk score and all available clinicopathologic parameters, including age, gender, Ann Arbor clinical stage, LDH ratio, ECOG performance status, number of extranodal site. Then we utilized the rms^39^ R packages set up a prognostic nomogram for OS probability assessment by enrolling all the independent prognostic factors. The discriminative efficacy, consistency and clinical judgment utility of the nomogram score was evaluated by time-dependent Calibration plots and decision curve analysis (DCA)^40^ using the rmda^41^ R package.

### Comprehensive analysis of molecular and immune characteristics in different IPI-IPM subgroups

DEGs between high and low risk score groups were analyzed following the fold change of a threshold at 1.5 and FDR value < 0.05. The gene expression of samples between IPI-IPM subgroups were analyzed with the t-distributed stochastic neighbor embedding (t-SNE) method by using the Rtsne^42^ package of R and then visualized on the 3D map with the scatterplot3d^43^ package of R. Somatic mutations of IPI-IPM subgroups were analyzed by using the Maftools^44^ R package. The intersection of DEGs and immune related gene set was used to construct a protein-protein interaction (PPI) network based on the STRING^45^ database.

The Cytoscape^46^ plugin MCODE^47^ and cytoHubba^48^ was utilized to identify top10 degree genes in the network. The TRRUST^49^ database was browsed to explore the curated transcriptional regulatory networks. Gene expression data (TPM) of 570 DLBCL samples were imported into CIBERSORT^50^, MCPCounter^51^ and xCell^52^ to calculate the score to estimate of the proportion of TME cells including immune and stromal cells.

### Clustering analysis of expression pattern of LME signatures

Unsupervised clustering algorithm was applied to analysis the gene expression pattern of LME signatures in 563 DLBCL samples. By using ConsensusClusterplus^53^ R package, we performed k-means clustering algorithm with 1000 repetitions to ensure the stability.

### Data Analysis

All statistical data was analyzed in the R software (version 4.0.3). A Wilcoxon test was applied to compare continuous variables between two groups of sample data. A Kruskal-Wallis test was applied to compare continuous variables among three or more groups of sample data. A test is considered with statistical significance at P < 0.05. Pearson correlation was used to test the linear correlation between two sets of data and an absolute Pearson correlation coefficient larger than 0.3 was considered to be correlated. A Pearson correlation is considered with statistical significance at FDR < 0.05. We used ggplot2, ggstatsplot and ggpubr R packages^54, 55^ for data analysis and visualization.

## Results

### Identification of immune related genes associated with IPI in DLBCL patients

A flowchart was diagramed to demonstrate the procedure and result of our study (**Figure 1**). RNA-seq data of 570 DLBCL patients were obtained from the TCGA (48 from TCGA-DLBC, 41 from CTSP-DLBCL1, 481 from NCICCR-DLBCL). Among the 566 DLBCL patients with survival (OS) data, 321 (56.71%) were male and 245 (43.29%) were female. Age of the patients at initial diagnosis ranged from 14 to 92 (median = 62). Other clinical characteristics including follow-up period, Ann Arbor stages, LDH ratio, ECOG performance status, and number of extranodal sites were all documented in **Table 1** and **Supplementary material 1**. To remove the batch effect among these 3 projects, we utilize the ComBat-seq function to transform the raw counts data with the sva R package. Then, the principal component analysis (PCA) was performed to show that there was no obvious batch effect among the samples (**Figure 2A**). After excluding samples with unrecorded IPI score or with IPI score crossing risk groups (such as 1-5 or 3-4), 458 DLBCL patients were divided into low-risk group (n = 118), intermediate-risk group (n = 221, 106 at low-intermediate risk and 92 at high-intermediate risk), and high-risk group (n = 109) (**Table 1**). Consistent with the previous publications, patients in the low-IPI risk group were shown with much longer overall survival (OS) (**Figure 2B**).

**Figure 1.**
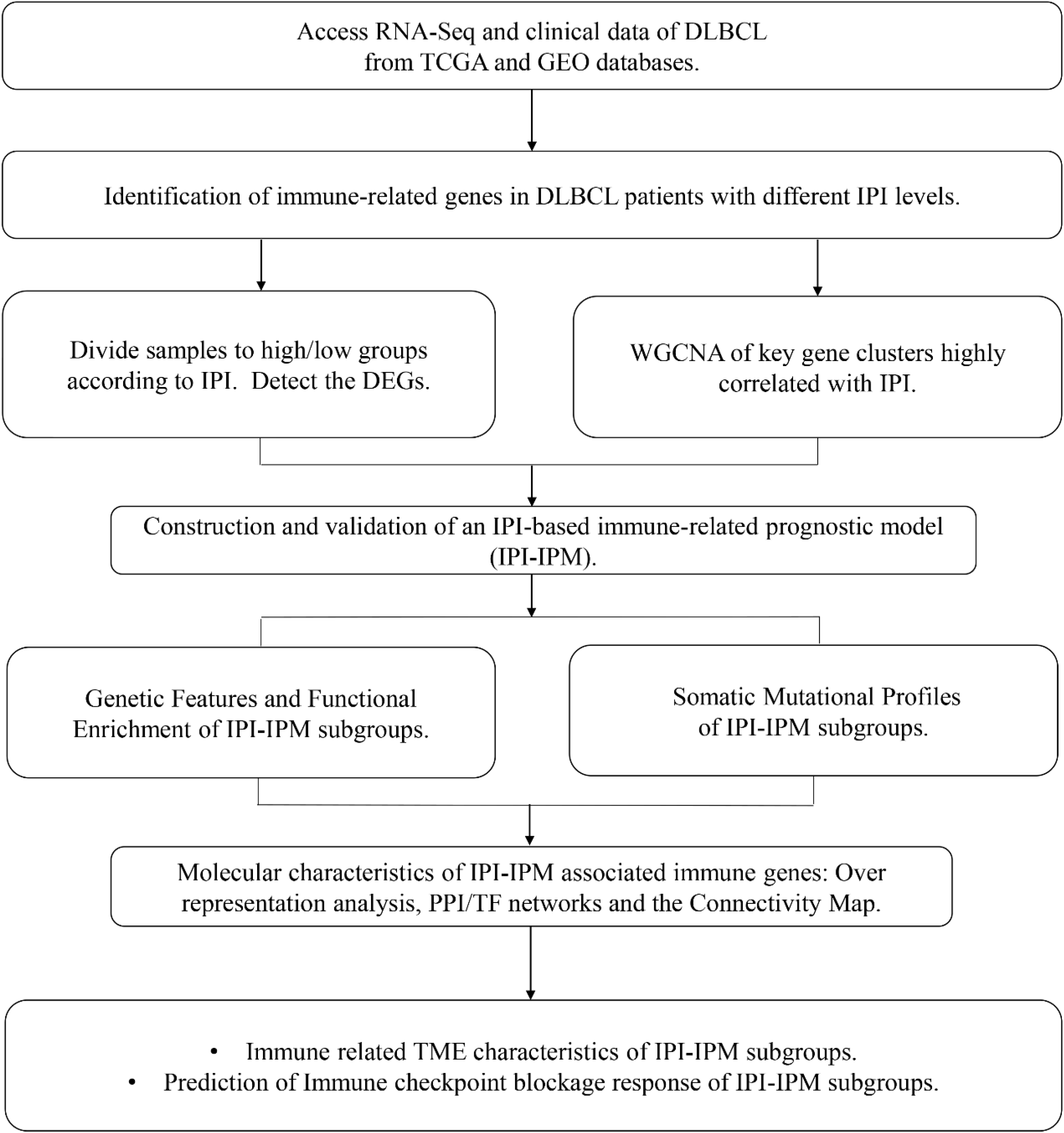
A flow chart for the process of the present study.

**Figure 2.**
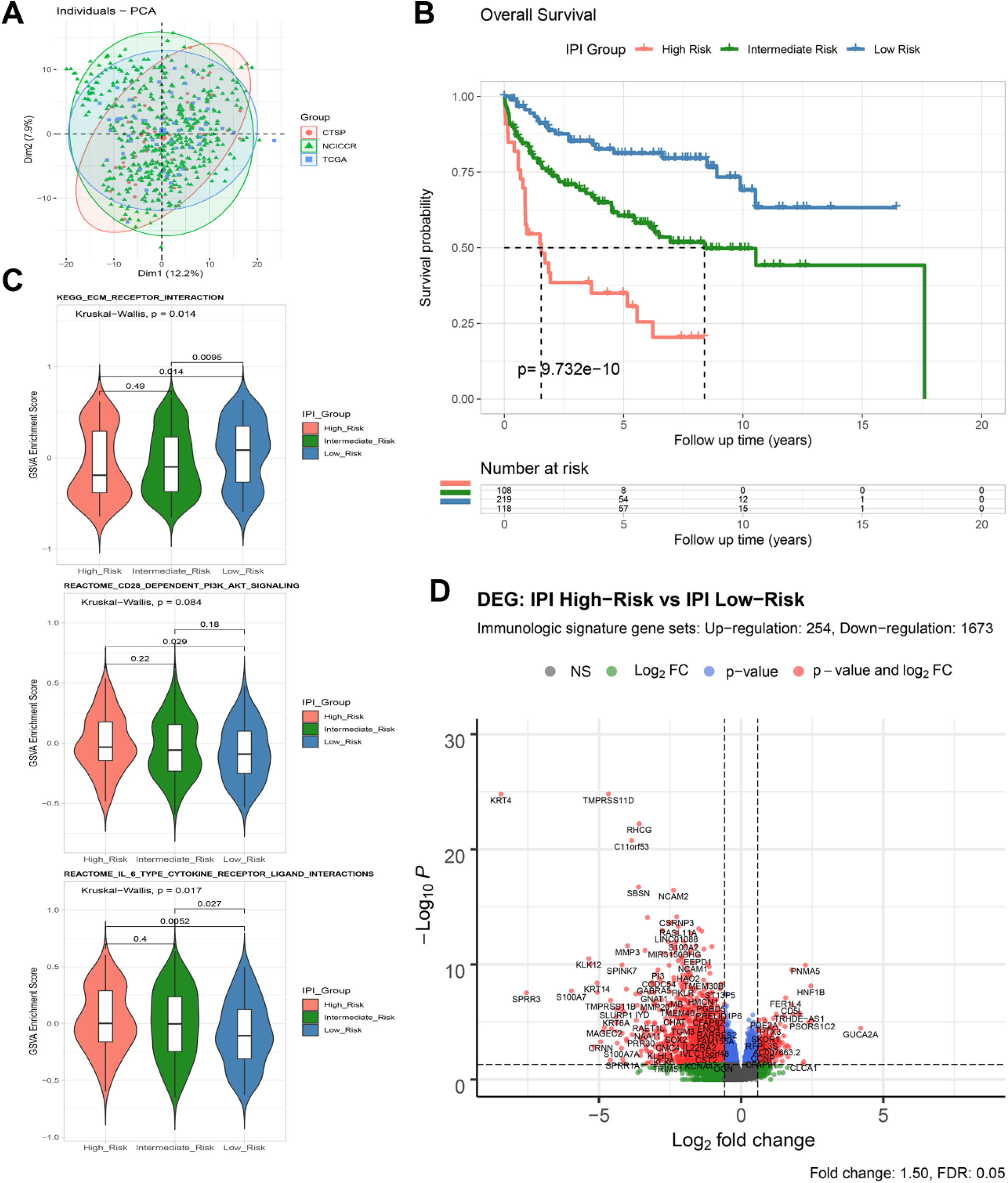
Identification of immune-related genes in DLBCL patients with different IPI levels. (A) PCA analysis. (B) Survival analysis of overall survival between high and low IPI groups. (C) Gene set variance analysis (GSVA) of enriched gene sets among different IPI groups. (D) Volcano plot of differentially expressed genes (DEGs) between the high and low IPI groups.

As shown in gene set variation analysis (GSVA), extracellular matrix (ECM) receptor interaction, CD28 dependent PI3K/AKT signaling and IL-6 type cytokine receptor ligand interaction and other immunologic signaling pathways were significantly enriched in low-risk group (**Figure 2C and Figure S1A-B**). A total 4651 genes (633 up-regulated and 4018 down-regulated) were detected significantly differentially expressed between the IPI high- and low-risk groups (**Figure S1C-D and Supplementary material 2**). By intersecting with the immunologic signature gene sets (combing 4686 genes from ImmuneSigDB and Immport, **Supplementary material 3**), 1927 immune DEGs (254 up-regulated and 1673 down-regulated) were identified for further analysis (**Figure 2D and S1E-F**).

We applied variance stabilizing transformed (VST) expression data via DESeq2 as the input data for WGCNA, including 13329 genes with the top 25% variance among all samples (**Figure 3A**). All clinical characteristics were enrolled as trait variables, and the best β value in the co-expression network was calculated to be 9 **(Figure S2B)**. Index for clustering of module eigengenes was modified to be 0.65, so as to construct reasonable number of merged modules **(Figure S2A-D)**. As shown in the Module-trait relationship, 8 modules were significantly correlated with IPI group (**Figure 3B**), and high correlation (P < 0.0001) between gene significance of immune score and gene module membership were found in the genes of 3 modules (Brown, Pink, Darkred) (**Figures 3C and S2E-G**). By intersecting 4106 genes from the top 3 IPI correlating modules with 1927 immune DEGs, a total of 371 genes were identified as immune related genes associated with IPI, which were used for prognostic risk model construction (**Figure S2H**).Several gene sets, such as ECM organization, ECM receptor interaction, PI3K/AKT signaling and integrin cell surface interaction, etc., were enriched in the top 3 IPI correlating modules (**Figure S2I and Supplementary material 4**).

**Figure 3.**
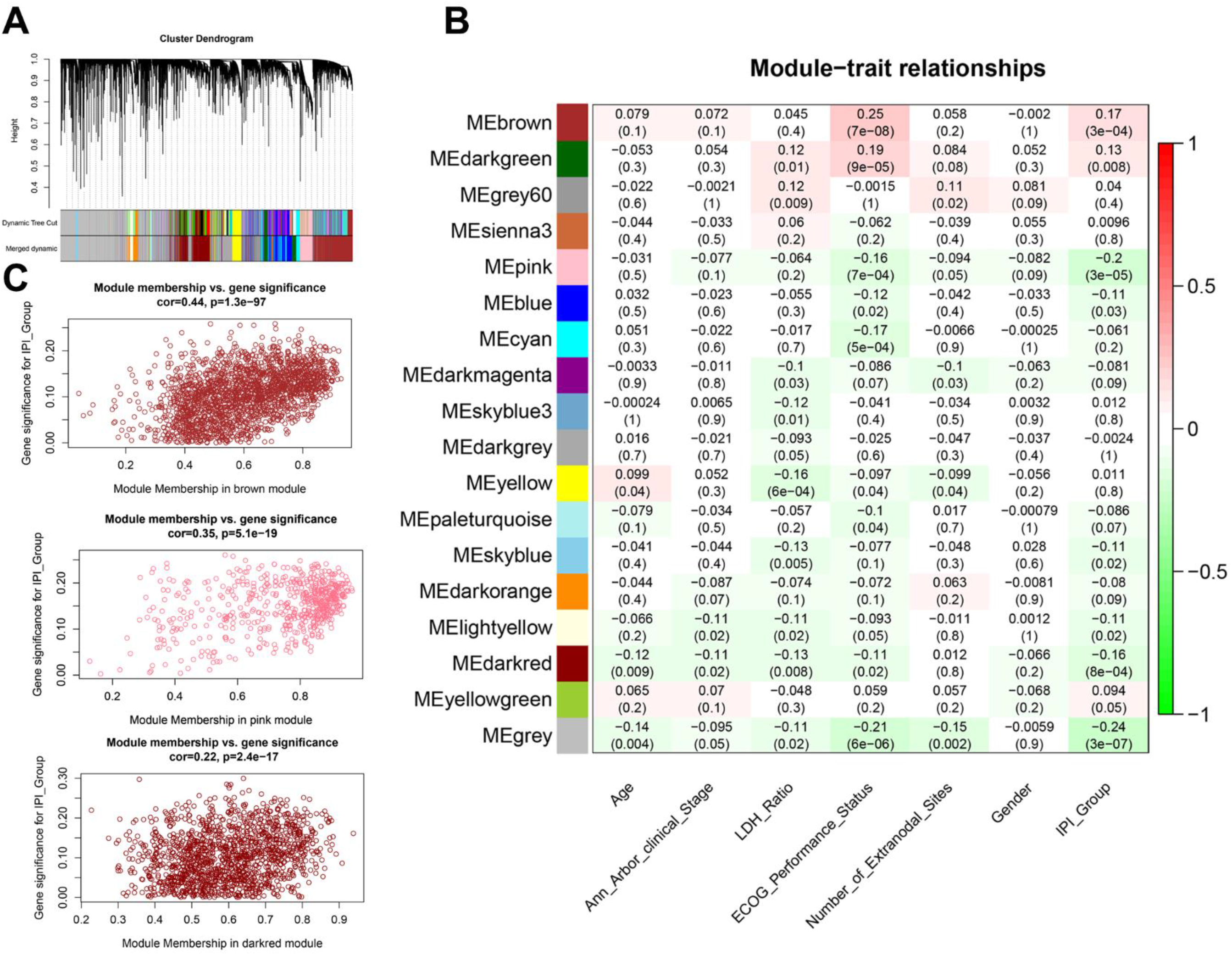
weighted gene co-expression network analysis (WGCNA) module identification and correlation analysis (A) Network analysis of gene expression in DLBCL identifies distinct modules of co-expression genes. (B) Analysis and visualization of Module-trait relationship to identify IPI related modules. (C) Correlation between gene module membership and gene significance for IPI in 3 modules.

### Construction and validation of an IPI-based immune prognostic risk model

In the training cohort (n = 563), 93 out of 371 IPI-related immune genes were significantly correlated with OS in the univariate Cox regression analysis (**Supplementary material 5**). Next, we applied the Lasso penalized Cox regression to identify the optimal number of genes (n = 10) for risk score model (**Figure 4A**). As a result of stepwise multivariate Cox regression and AIC analysis, 8 genes were selected to construct the most optimal IPI-based immune prognostic model (IPI-IPM) (**Figure S3A-B and 4B**). Risk score = (expression level of CMBL * 0.360 + expression level of TLCD3B * (−0.350) + expression level of SYNDIG1 * (−0.247) + expression level of ESM1 * (−0.238) + expression level of EPHA3 * (−0.163) + expression level of HUNK * (−0.156)+ expression level of PTX3 * 0.138 + expression level of IL12A * 0.111). As shown in the time-dependent ROC curves, AUCs were 0.703, 0.738, 0.733 for the 1, 3, 5-year, respectively (**Figure 4C**). The C-index of the risk score model was 0.732 (95% CI: 0.684-0.779, P = 1.38e-21). These results showed that IPI-IPM endowed a good capacity in OS prediction. According to the ROC curve of the median survival time (**Figure S3C**), we identified 0.982 as the cut-off value. Then we calculated the risk score of each patient and divided them into high- and low-risk groups. As shown in **Figure 4D**, Kaplan-Meier survival analysis showed shorter OS of patients in the high-risk score group (log-rank P = 3.13e-14). The distribution of the risk score, survival status and the 8-gene expression between high- and low-risk score groups was shown in **Figure S3D and Figure 4E**.

**Figure 4.**
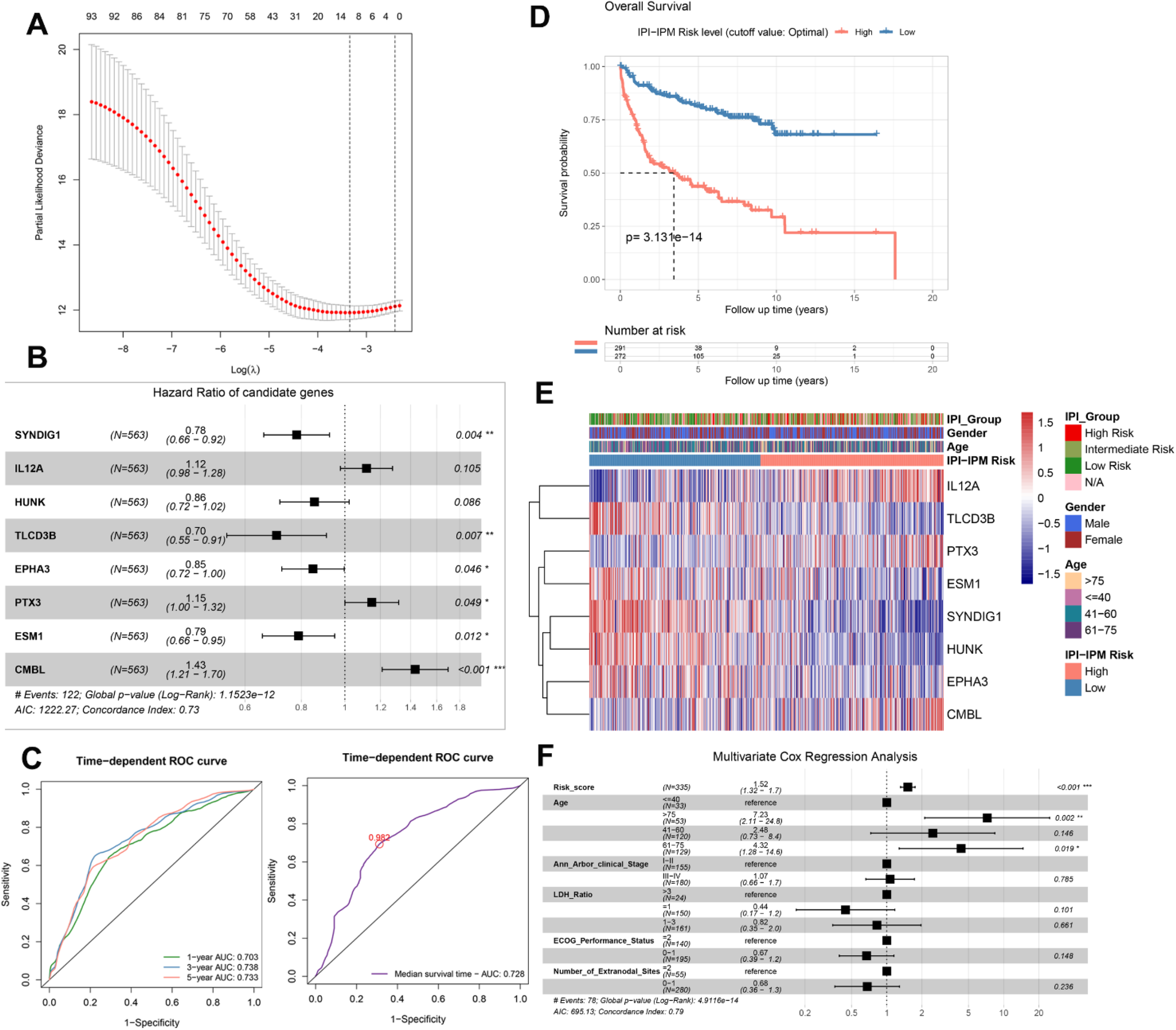
Construction of an IPI-based immune prognostic model. (A) Plot of the Lasso penalized Cox regression. (B) A forest plot of the genes composing the IPI-based immune prognostic model (IPI-IPM). (C) time-dependent ROC curves for the IPI-IPM. (D) Survival analysis of overall survival between high and low IPI-IPM risk groups. (E) Heatmap of the gene expression in high and low IPI-IPM risk groups. (F) The multivariate analysis of IPI-IPM risk score and clinicopathologic parameters including Age, Ann Arbor clinical stage, LDH ratio, ECOG performance status and the number of extranodal sites.

Moreover, 335 patients with all available clinicopathologic parameters were enrolled for Multivariate Cox regression analysis, presenting the risk scores and age as independent prognostic factors of OS (**Figure 4F**). Risk scores along with age, Ann Arbor clinical stage, LDH ratio, ECOG performance status and number of extranodal sites were integrated to construct a nomogram model, showing IPI-IPM risk scores with the highest risk points (ranging from 0 to 100, **Figure 5A**). As shown in the DCA analysis, the nomogram along with risk score from IPI-IPM showed relatively high net benefit (**Figure 5B**). Moreover, the bias-corrected lines for the nomogram were shown close to the ideal line in 1,3,5-year and the median survival time periods (**Figure 5C**). The C-index for the nomogram was 0.790 (95% CI: 0.736-0.843, P = 2.38e-26). Altogether, these results suggested that the nomogram had excellent capacity and consistency for OS prediction in the training cohort.

414 patients from GEO (GSE10846) with survival data and microarray data were enrolled for further validation of IPI-IPM. The risk score for each patient was calculated and all patients were divided into high- and low-risk groups likewise. As shown in **Figure 5D**, AUCs were 0.619, 0.603, 0.601 for the 1, 3, 5-year, respectively. Kaplan-Meier survival analysis showed significantly shorter OS of patients in the high-risk group (log-rank P = 5.30e-05, **Figure 5E and S3E-F**). Moreover, patients in the high-risk group were shown to have shorter OS no matter which treatment regimens they received (**Figure 5F**). Taken the results of training and testing cohorts together, the IPI-IPM and the nomogram combining risk score with relevant clinical characteristics (age, Ann Arbor clinical stage, LDH ratio, ECOG performance and number of extranodal site) was an excellent model for predicting short-term or long-term OS in DLBCL patients, which might guide the therapeutic strategy decision and long-term prognosis follow-up.

**Figure 5.**
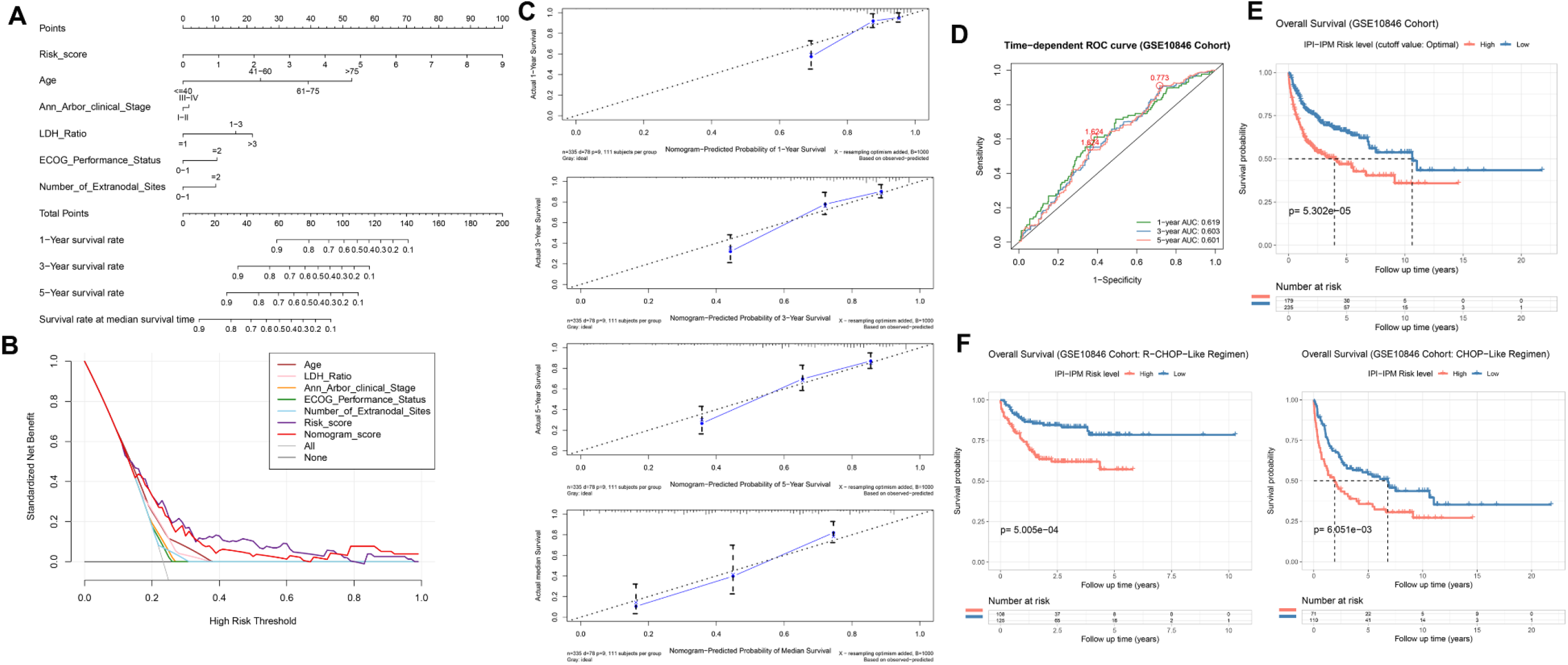
Validation of the IPI-IPM and Construction of an IPI-based immune nomogram model. (A) Nomogram for the prediction of the survival probability of 1-, 3-, and 5-year overall survival. (B) The DCA analysis of all parameters in the nomogram. (C) Calibration plots of nomogram-predicted probability of 1-, 3-, 5-year, and median survival. (D) time-dependent ROC curves for the Nomogram. (E-F) Survival analysis of overall survival between high and low risk groups in the testing (GSE10846) cohort, in patients with R-CHOP like regimens and those with CHOP like regimens (F).

### Molecular characteristics of IPI-IPM subgroups

Compared to the low-risk group, A total 5980 genes (690 up-regulated and 5290 down-regulated), and 2731 immunologic genes (400 up-regulated and 2331 down-regulated) were detected significantly differentially expressed in high-risk group (**Figure S4A and 6A, Supplementary material 6**). The pre-ranked GSEA was performed to display that several gene sets including negative regulation of immune response, DNA repair and response to IL-12, etc., were enriched in high-risk group, and several gene sets including ECM receptor interaction, IL-10 synthesis and regulation of humoral immune response, etc., were enriched in low-risk group. Details were all documented in **Figure 6B and S4B-E** and **Supplementary material 6**. Additionally, t-SNE was applied to show an obvious genetic diversity between high- and low-risk groups (**Figure 6C**). Based on the Pearson correlation analysis, there existed either positive or negative correlation between these 8 genes from IPI-IPM (**Figure 6D)**. And **Figure 6E** presented the heatmap of differentially expressed genes correlating IPI-IPM risk scores between high- and low-risk groups.

**Figure 6.**
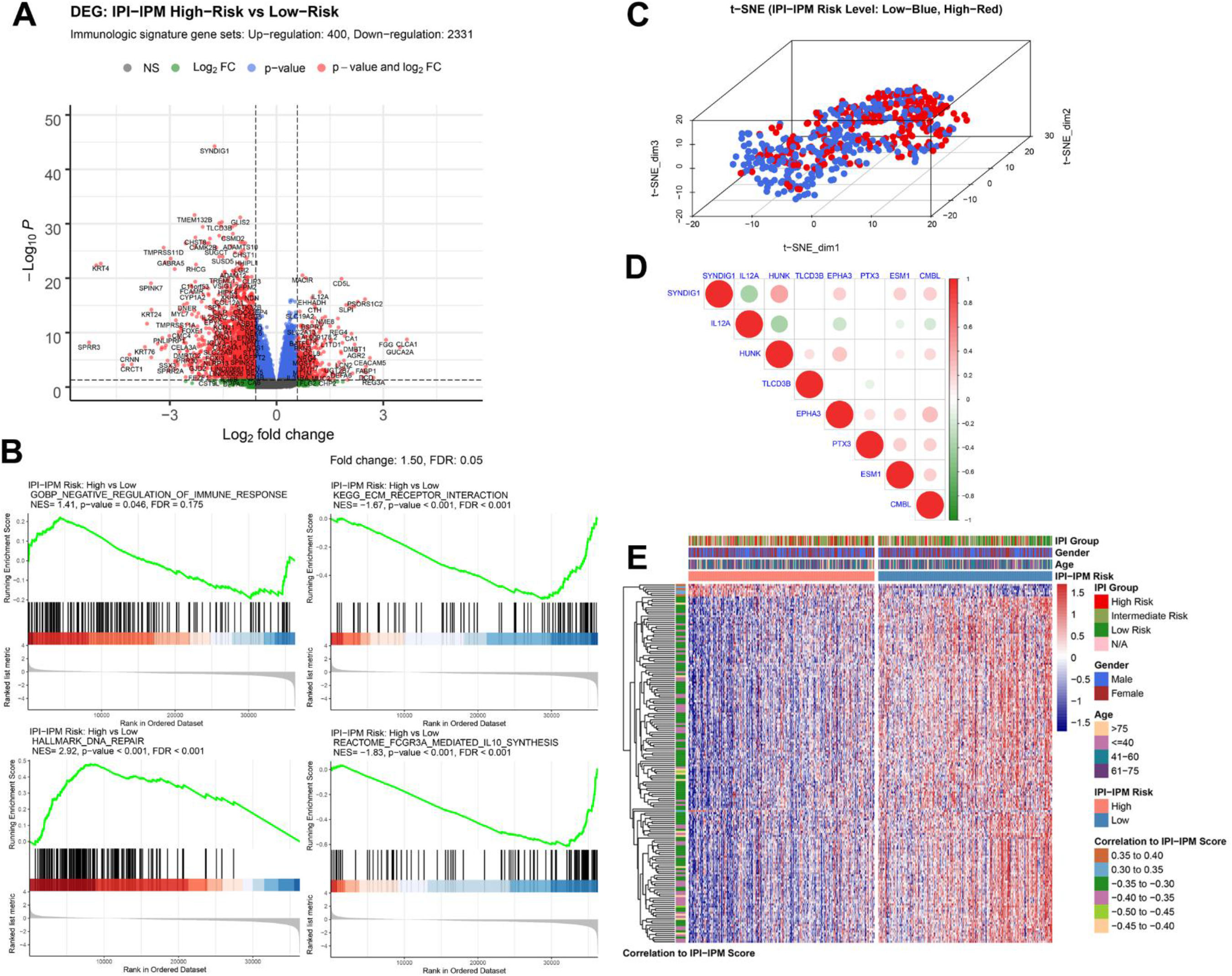
Molecular characteristics of IPI-IPM associated immune genes. (A) Volcano plot of immune related DEGs between high and low IPI-IPM risk group. (B) Preranked GSEA of enriched gene sets between high and low IPI-IPM risk group. (C) The t-SNE algorithm were applied to show the difference of DLBCL patients between the high and low IPI-IPM risk groups. (D) Pearson correlation analysis of the 8 genes composing the IPI-IPM. (E) Heatmap of immune related genes correlating the IPI-IPM risk groups.

To gain further molecular insight into the molecular characteristics of IPI-IPM, 176 genes correlating with risk scores and these 8 genes from IPI-IPM were identified as IPI-IPM associated immune genes (absolute Pearson correlation coefficient ≥ 0.3, FDR < 0.05). Over representation analysis was applied to identify the enriched biological functions and pathways, such as ECM organization and T cell differentiation, activation and mediated immunity (**Figure 7A-B**). Detailed results were listed in **Supplementary material 6**. As shown in the PPI network, ECM organization, oncogenesis and tumor immunity associated regulatory genes (COL6A2, COL16A1, COL26A1, COL13A1, COL22A1, C3, ELN, MMP9, CLU, FOXP3,ADGRL1) were closely correlated, acting as hub genes (**Figure 7C**). In addition, TFTRUST database was applied to explore the transcription factors (TFs) regulating the 184 IPI-IPM associated immune genes, thus MMP9, FOXP3 and PLAU were identified in the TF network (**Figure 7D**).

**Figure 7.**
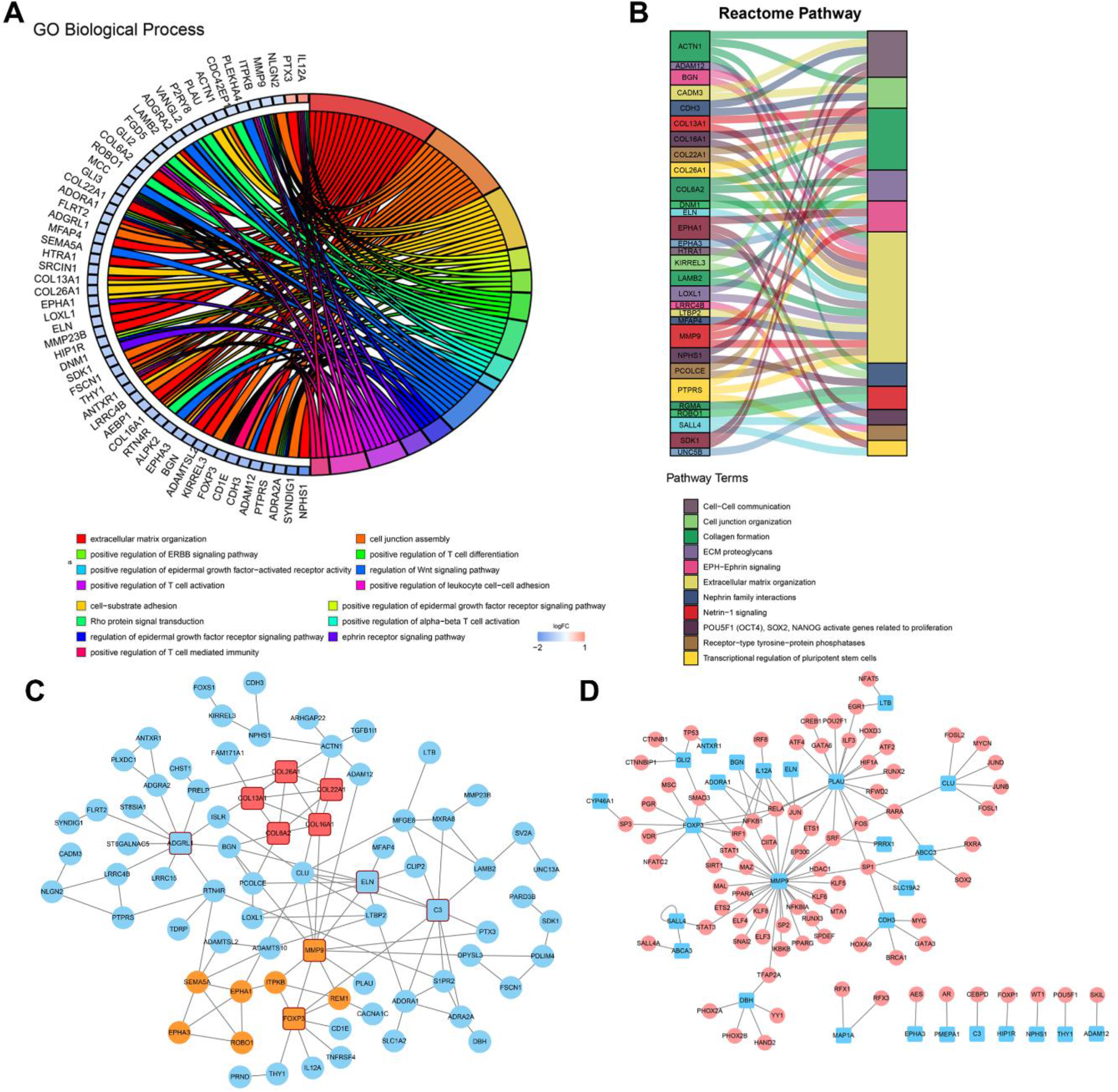
Immune characteristics of IPI-IPM subgroups. (A and B) Over representative analysis: A Chord map of the enriched Biological processes (A) and a Sankey plot of the enriched Reactome pathways (B). (C) Protein-protein interaction network (PPI) based on the STRING database. (D) Transcription factor network of immune related genes correlating the IPI-IPM risk groups.

Based on the somatic mutational data of 37 samples from TCGA, 18 out of 20 patients in high-risk group and 13 out of 17 patients in low-risk groups were found with altered gene expression (**Figure S5A-B and 8A**). Although most mutations were missense mutation, more nonsense and other mutations were identified in high-risk group (**Figure 8A**). Besides, the mutation frequency of top10 genes in high-risk group was much higher than that in low-risk group. Furthermore, we investigated specific mutation sites of key genes corresponding to their amino acids location, including KMT2D, MUC16, CARD11, LRP1B, BTG2 and PIM1, etc. (**Figure 8B and S5C**). As shown in the Oncodrive plot, MyD88, CD79B, KHL6 and MUC4 were identified as cancer driver genes in high-risk group while only PEG3 and ZNF337 were identified in low-risk group (**Figure 8C and Supplementary material 7**).

**Figure 8.**
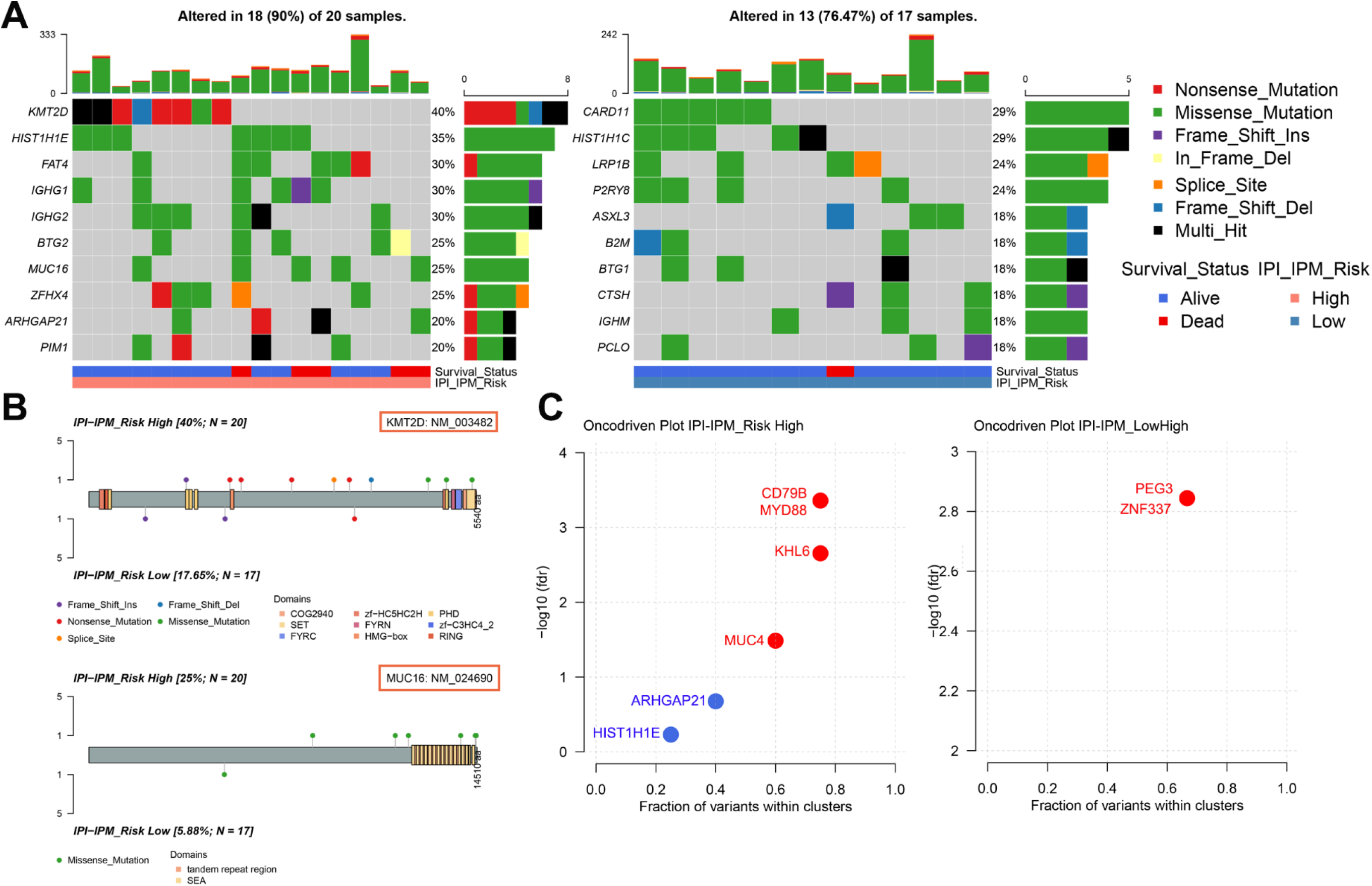
Somatic mutational profiles of IPI-IPM subgroups. (A) Differentially mutated genes between high and low IPI-IPM risk groups. Top 10 Mutated genes (rows) are ordered by mutation rate. The color-coding legends indicates the mutation types and survival status of patients. (B) Lollipop plots for amino acid changes of KMT2D and MUC16. (C) Oncodrive plots of high and low risk IPI-IPM groups.

### Immune landscape of IPI-IPM subgroups

CIBERSORT was applied to analyze the infiltrating abundances of various immune cell types in different IPI-IPM subgroups (**Figure 9A and S6A**). Activated memory CD4^+^ T cells and resting NK cells were highly infiltrated in high-risk group, while memory B cells, CD8^+^ T cells, follicular helper T cells, regulatory T cells (Tregs), and non-activated macrophages (M0) were more abundant in low-risk group (**Figure 9B**). In addition, MCPCounter and xCell algorithm were applied to show that myeloid dendritic cells (mDCs) and common lymphoid progenitors were highly infiltrated in high-risk group, while hematopoietic stem cells (HSCs), cancer associated fibroblasts (CAFs) and T cells especially CD8^+^ T cells were more abundant in low-risk group (**Figure 9C and S6B**).

**Figure 9.**
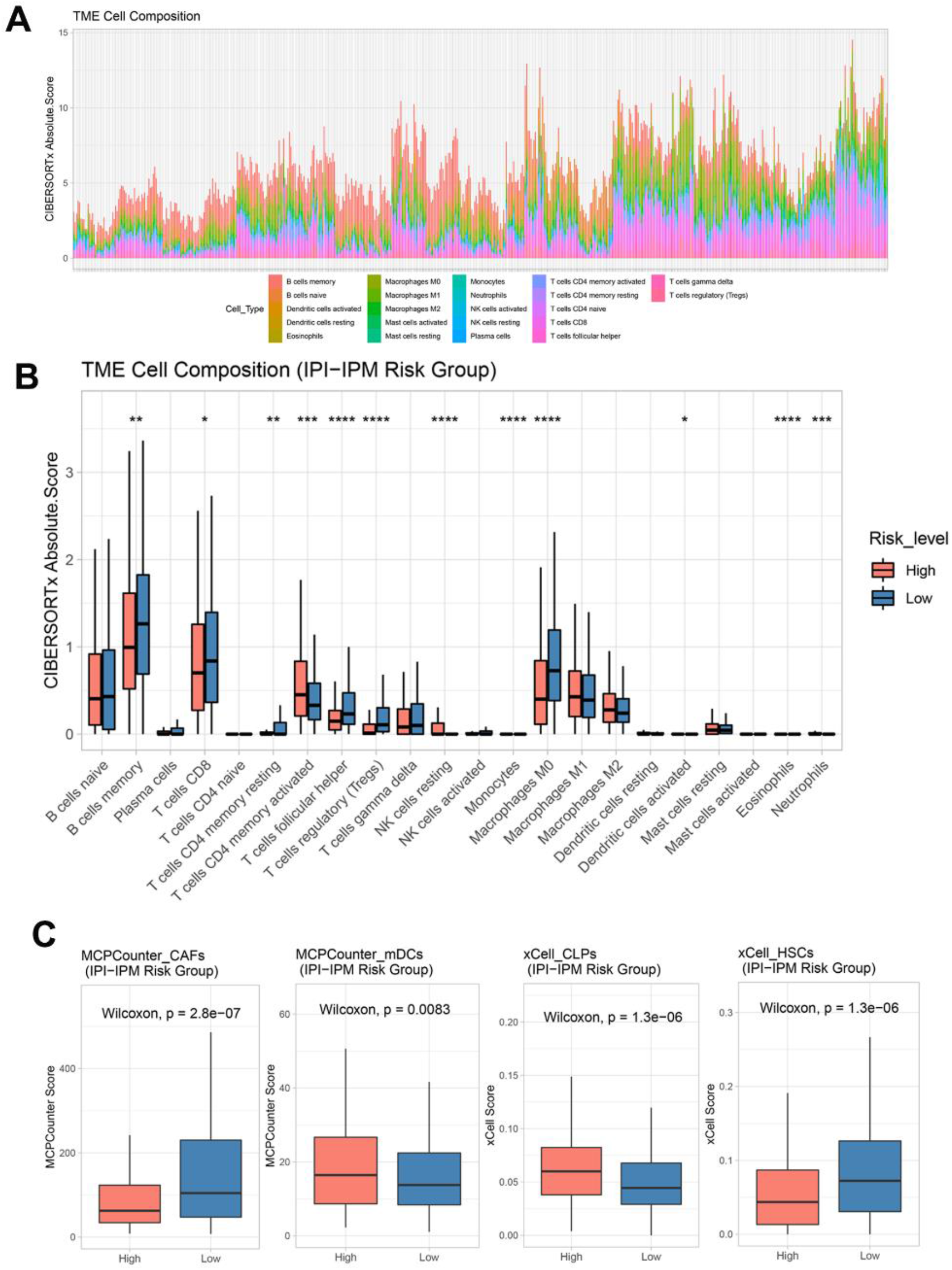
Tumor immune microenvironment (TME) characteristics of IPI-IPM subgroups. (A-B) Analysis of immune cells infiltration by using the CIBERSORT algorithm: Relative proportion of each type of cells infiltration in DLBCL patients (A) and a bar plot visualizing significantly differentially infiltrated cells between high and low IPI-IPM risk groups (B). (C) Analysis of immune cells infiltration by using the MCPcounter and xCell algorithm.

In addition, DLBCL subtypes varied between high-and low-risk groups. As shown in **Figure 10A**, more patients with higher IPI level were classified into IPI-IPM high-risk group, with similar results about age, Ann Arbor clinical stage, LDH ratio, ECOG performance and the number of extranodal sites (**Figure S6C**). Whereas high-risk group contained a higher proportion of ABC-DLBCLs, low-risk group was more enriched in GCB-DLBCLs (P = 1.34e-14). Schmitz et al.^4^ identified four prominent genetic subtypes in DLBCL with different responses to immunochemotherapy, termed MCD (the co-occurrence of MYD88^L265P^ and CD79B mutations), BN2 (BCL6 fusions and NOTCH2 mutations), N1 (NOTCH1 mutations), and EZB (EZH2 mutations and BCL2 translocations), and uneven distribution of these 4 subtypes were found between high- and low-risk groups. For example, the poor-prognostic MCD subtype represented 40% of high-risk group and 12% of low-risk group. Also, the correlation analysis showed IPI-IPM risk scores positively correlated with the expression Bcl-2 and Myc, and negatively correlated with Bcl-6 expression (**Figure 10B**). Taken together these data indicates that IPI-IPM provide additional orthogonal information from previous lymphoma classifications.

**Figure 10.**
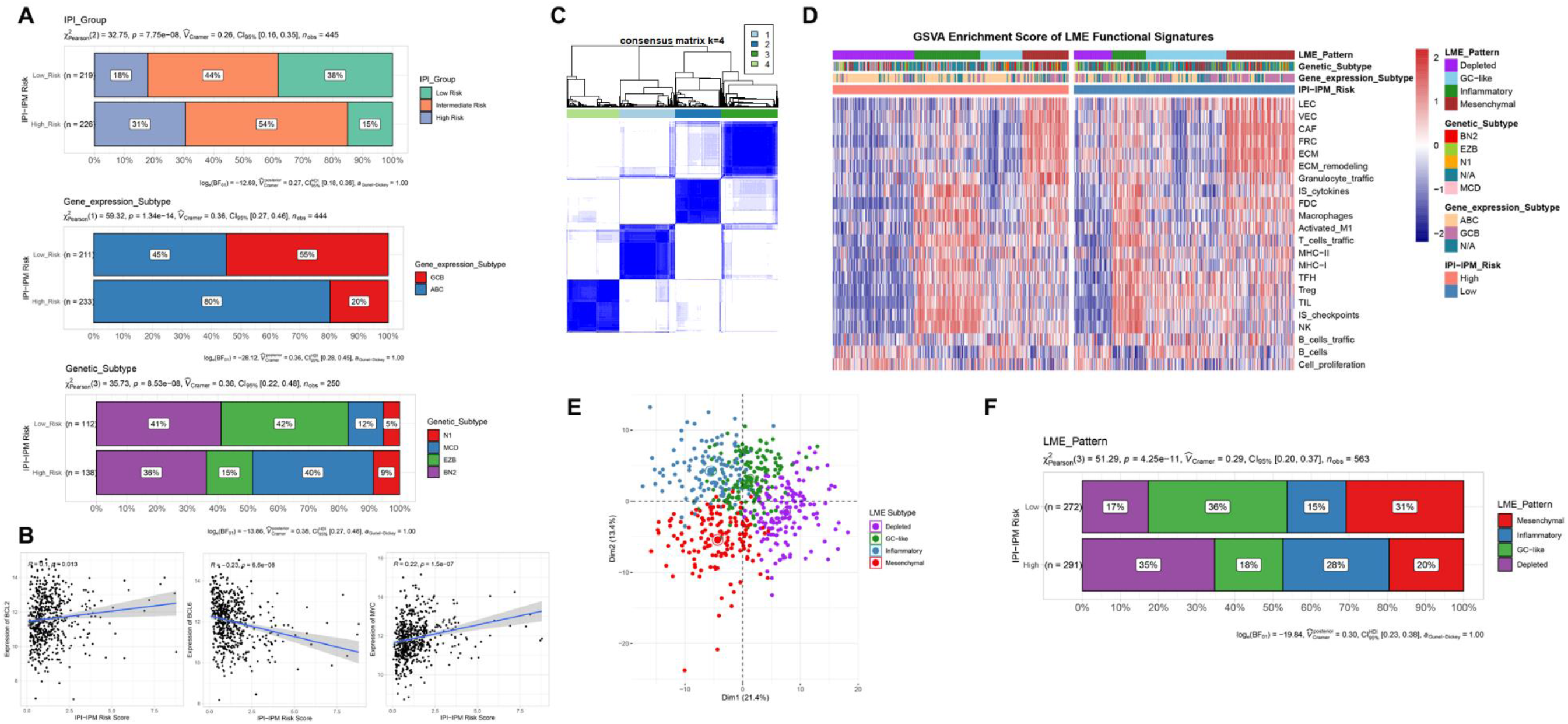
Molecular and microenvironmental subtypes of DLBCL. (A) Distribution of IPI groups, gene expression subtypes and genetic subtypes between high and low IPI-IPM risk groups. (B) Correlation analysis of IPI-IPM risk score and Bcl-2, Bcl-6 and c-Myc. (C) Consensus clustering and communities detection of lymphoma microenvironment (LME) clusters. (D-E) Distribution of LME patterns between high and low IPI-IPM risk groups. (F) GSVA enrichment score of LME functional signature between high and low IPI-IPM risk groups.

To further explore the interaction between DLBCL cells and the microenvironment, Kotlov et al. ^11^defined the lymphoma microenvironment (LME) into 4 major transcriptionally-defined categories with distinct biological properties and clinical behavior, including “germinal center-like” (GC), “mesenchymal” (MS), “inflammatory” (IN) and “depleted” (DP) form, respectively. Similarly, we utilized an unsupervised clustering method to assign the samples into 4 groups by using expression data of the 22 functional gene expression signatures (F^GES^) sets (**Figure 10C**). GSVA enrichment scores were calculated to demonstrate the distinct TME characteristics among the 4 biological patterns (**Figure 10D**). Consistent with CIBERSORT results, the DP LME categories represented 35% of high-risk group whereas GC-like and MS LME categories were more enriched in low-risk group (**Figure 10E-F and Supplementary material 8**).

### Potential therapeutic value of IPI-IPM

To further understand the effects of the risk score on drug response, 184 IPI-IPM associated immune genes mapped into the Connectivity Map database^56^. As shown in **Figure 11A and S7**, 14 genes (ADORA1, ADRA2A, CACNA1C, DBH, DNM1, ELN, ENPP1, EPHA1, IL12A, MMP9, PLAU, S1PR2, SLC1A2 and SV2A) were associated with 122 inhibitors, involved with 54 mechanisms of action (MOA). Details were documented in **Supplementary material 9**.

**Figure 11.**
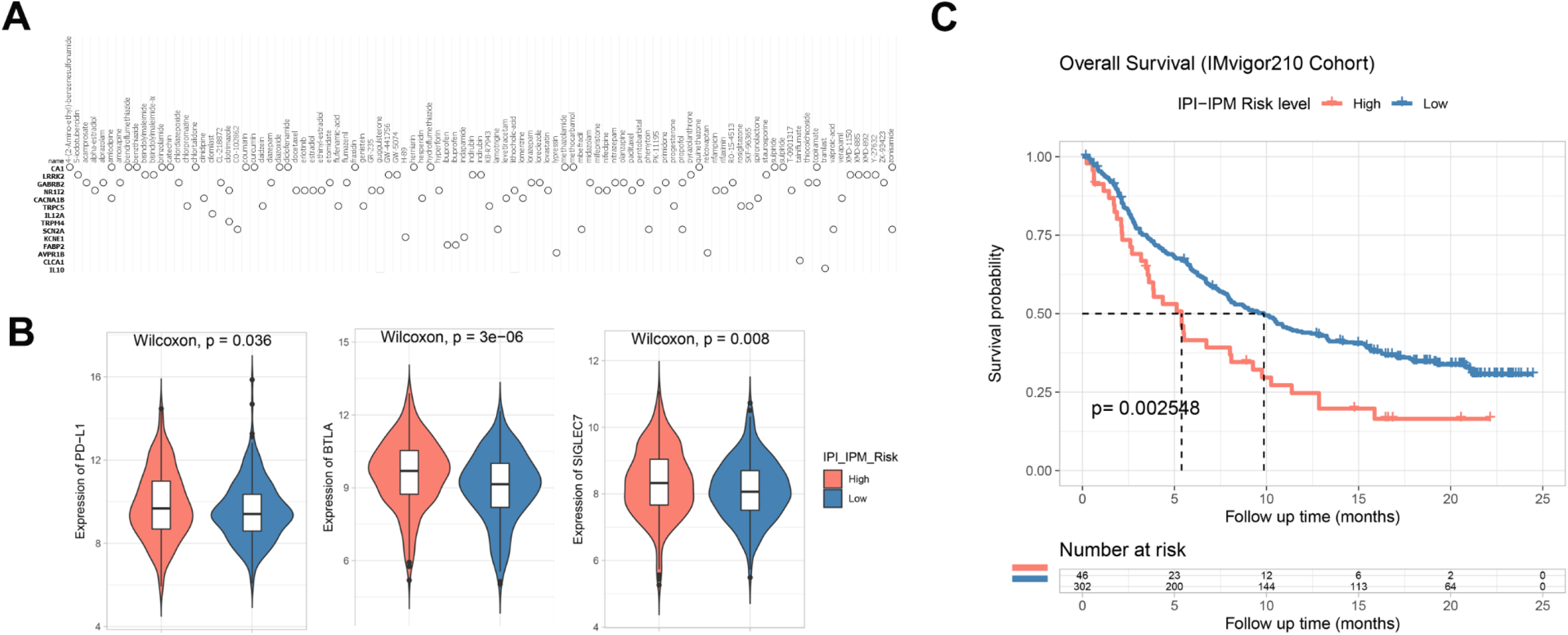
Potential therapeutic value of IPI-IPM. (A) Connectivity map of top enriched immune related genes correlating the IPI-IPM risk groups. (B) The expression of inhibitory immune checkpoints between high and low IPI-IPM risk groups. (C) Survival analysis of IMvigor210 Cohort.

The clinical development of cancer immunotherapies, along with advances in genomic analysis, has validated the important role of TME in predicting cancer response to immune checkpoint blockade therapy (ICB)^13, 57, 58^. We investigate the expression of several inhibitory immune checkpoints between high- and low-risk groups using TMM normalized counts data. As shown in **Figure 11B**, the expression of PD-L1, BTLA and SIGLEC7 were significantly up-regulated in high-risk group. Details were documented in **Supplementary table 10**.

Since various biomarkers were reported to predict the response to immunotherapy, including tumor mutation burden and immune checkpoints expression such as PD-L1. We examined the value of the IPI-IPM to predict the response of patients to ICB therapy, based on a publicly accessible dataset - IMvigor210 Cohort^26^. Using the same cut-off point for group assigning, we observed that patients with high IPI-IPM risk had shorter OS after anti-PD-L1 treatment (log-rank P = 0.003, **Figure11C**).

## Discussion

Recent groundbreaking insights into the pronounced genomic heterogeneity of DLBCL have identified potential biomarkers for patient diagnosis and prognosis, paving the way for a standardized application of precise medicine^3, 5, 13^. Multiple subtype classifications as well as IPI and enhanced NCCN-IPI were built to stratify prognostically relevant subgroups of DLBCL patients with R-CHOP therapy, the robustness of which however needs to be further investigated in the context of targeted therapies^7, 11, 13, 59^. In the current study, we used WGCNA to profile IPI correlating immune gene sets, constructed and validated an 8-gene IPI-IPM (CMBL, TLCD3B, SYNDIG1, ESM1, EPHA3, HUNK, PTX3 and IL12A), with shorter OS in high-risk patients and longer OS in low-risk patients in both TCGA and GEO cohorts.

ESM1, also known as endocan, was initially proven to regulate endothelial cell function, also participate in the initiation and progression of human cancers including esophageal cancer, hepatocellular carcinoma, bladder cancer and breast cancer^60^. EphA3 receptor plays critical role in cell adhesion and migration during development and homeostasis of many tissues, as well as cancer growth, progression and angiogenesis^61^. Also, EphA3 high expression in tumors but not in normal tissues, together with antitumor properties of anti-EphA3mAb (chIIIA4), defined EphA3 as a potential target for antibody-based anticancer therapies^61^. PTX3 is secreted by dendritic cells, macrophages, and fibroblasts, and actively involved in regulation of inflammation, tissue remodeling and cancer^62^. PTX3 interacts with the PI3K/AKT/mTOR signaling pathway or the fibroblast growth factor-2 (FGF2) / FGF receptor (FGFR) system to regulate tumor cell proliferation, apoptosis and metastasis in lung cancer, breast cancer, melanoma, prostate cancer and multiple myeloma^62, 63^. IL12A is a potent immune suppressive cytokine produced by regulatory B (Breg) cells, Treg cells, macrophages, dendritic cells and tumor cells, which suppresses the effector functions of CD4^+^ and CD8^+^ T cells but strongly favors Tregs proliferation^64^. Furthermore, Larousserie et al. found that high levels of IL12A were associated with the poor survival of DLBCL patients^65^.

High throughput gene expression databases and bioinformatic analysis have enabled systematic profiling of prognostic signatures in DLBCL. For example, Zhou et al. uncovered differentiated lncRNA expression pattern between GCB and ABC DLBCL and identified an immune-associated 17-lncRNA signature for subtype classification and prognosis prediction^66^. Chapuy et al. defined five distinct DLBCL subsets by integrating recurrent mutations, SCNAs and SVs^5^. And Hu et al. built a predictive model comprising drug resistance signature along with clinical factors including age at diagnosis, stage, the number of extra nodal sites and the ECOG performance score^59^. In the current study IPI-IPM risk score remained an independent prognostic factor after modification of clinical characteristics, thus we developed a nomogram model combining the risk score and other clinical features (age, Ann Arbor clinical stage, LDH ratio, ECOG performance and number of extranodal site) to predict OS probability of DLBCL patients in 1-, 3-, 5-year and the median survival time, respectively. Both the calibration curve and DCA analysis supported that our nomogram provides a complementary perspective on individualizing tumors and develops an individual scoring system for patients, thus arising to be a promising tool for clinicians in the future.

In the GSEA analysis between high- and low-risk groups, several immune-related gene sets, including negative regulation of immune response, B cell and T cell proliferation, response to IL-12 and γ-IFN, and NOD-like and TOLL-like receptor signaling, were enriched in high-risk group whereas ECM receptor interaction, IL-10 synthesis and regulation of humoral immune response, etc., were enriched in low-risk group. Therefore, we speculated that the local immune signature conferred a weaker immune phenotype in high-risk group but an intense immune phenotype in low-risk group. Moreover, over representation analysis identified several immune-related pathways including ECM organization and T cell activation were enriched with IPI-IPM associated immune genes. ECM has important roles in supporting the cells and regulating intercellular interactions, thus contributing to progression of several malignancies^11, 14^. Lenz et al. built a survival model with 2 stromal gene signatures for DLBCL patients who received CHOP or R-CHOP, where the prognostically favorable stromal-1 signature reflected ECM deposition and histiocytic infiltration while the prognostically unfavorable stromal-2 signature reflected tumor blood-vessel density^12^.

To further explore the immunological nature of the IPI-IPM subgroups, we analyzed 37 samples’ somatic mutational profile and found higher mutation counts in high-risk group with more nonsense mutations, although missense mutations were the most common type. And the largest difference in mutations between high- and low-risk groups was in KMT2D mutation (40% in high-risk samples vs. 17.65% in low-risk samples)^67^. KMT2D is a tumor suppressor gene in DLBCL and genetic ablation of KMT2D in a BCL2-overexpression driven model promotes a higher penetrance of DLBCL^68^. Moreover, MyD88 and CD79B were identified as cancer driver genes in high-risk group, in good agreement with the previous findings that MYD88 and CD79B mutations have been associated with tumor response and survival in DLBCL patients^69-71^. Ngo et al. initially identified activating MyD88 mutations in DLBCL, where L265P was the most frequent and oncogenic form. MyD88 was shown to interact with IL-1 receptor-associated kinase (IRAK) 1 and IRAK4, activate the NF-κB and JAK-STAT3 pathways, promoting malignant cells proliferation and causing worse survival of DLBCL patients^72^. CD79B mutations frequently are detected in the first tyrosine (Y196) of the immunoreceptor tyrosine-based activation motif (ITAM). CD79B mutations were shown to cause chronic active BCR signaling and constitutive NF-κB activation, further promoting tumor cells growth within immunosuppressive TME.

Consistently, the composition of immune cells was different between two IPI-IPM subgroups, where memory B cells, CD8^+^ T cells, Tregs, M0 and CAFs were more abundant in low-risk group. It’s generally accepted that cytotoxic CD8^+^ T cells, following successful priming, recognize tumor-specific (neoantigens) or tumor-associated antigens and exert anti-tumor function primarily via the release of cytotoxic molecules such as perforin and granzymes^73-75^. However, Tregs suppress CD8^+^ T cells by direct cell contact and secretion of inhibitory cytokines including IL-10 and TGF-β^74, 76^. TAMs have been shown to mediate antibody-dependent cellular phagocytosis of rituximab engaged malignant B cells, but limit CD8^+^ T cells activity through PD-L1 expression, as well as releasing IL-10 and TGF-β or inhibiting enzymes, thus regulating anti-tumor immunity and response to therapy^77^. CAFs, the resident fibroblasts activated in a chronic inflamed TME, have been shown to promotes recruitment and polarization of regulatory cells including Tregs, monocytes and M2 by secreting IL-6, CXCL12, Chi3L1, MCP-1 and SDF-1, thus actively shaping the immune infiltration in TME^78^. In addition, CAFs have been shown to impact the cytolytic activity of CD8^+^ T cells through different mechanisms, such as producing prostaglandin E2 (PGE2) and NO to dampen CD8^+^ T cells proliferation, expressing PD-L2 and FasL to promote CD8^+^ T cells apoptosis, and inducing abnormal ECM deposition and remodeling to physically trap CD8^+^ T cells and prevent effective tumor access^79^.

Besides, DP LME represented more of high-risk group while immune rich GC LME was more enriched in low-risk group. Kotlov et al. characterized the microenvironment of DLBCL by analyzing gene expression profiles, developing TME-derived F^GES^, analyzing ECM composition by proteomics and establishing patient-derived tumor xenografts (PDTX) models^11^. Four basic categories of DLBCL LME with distinct clinical and biological connotations were identified to uncover the bidirectional interaction between DLBCL cells and LME. Remarkably, immune rich GC LME confers a better prognosis than DP LME, suggesting the fundamental role of LME in preventing lymphomagenesis. In turn, DLBCL cells develop genetic and epigenetic traits that contribute to immune evasion from LME.

Finally, we applied Spearman correlation analysis to estimate the potential therapeutic effects of IPI-IPM; and explored the expression of several inhibitory immune checkpoints between IPI-IPM subgroups to predict the response to immunotherapy. The results shown that IPI-IPM associating genes were correlated with sensitivity to drugs targeting Aurora kinase, DNA methyltransferase (DNMT), Histone acetyltransferase (HAT), FLT3, EGFR and VEGFR signaling pathways, indicating that high-risk DLBCL patients may benefit from novel inhibitors targeting these signaling pathways. As for the inhibitory checkpoints expression, PD-L1, BTLA and SIGLEC7 were significantly up-regulated in high-risk group. Upregulation of checkpoints (PD-1, CTLA-4, and TIM-3) and their ligands (PD-L1 and PD-L2) in TME can mediate tumor cells to escape immune surveillance by modulating T-cell activity^13, 57, 80, 81^. Thus, ICB has exerted significant antitumor effect in both solid tumors and hematologic malignancies. Although nivolumab monotherapy showed a low efficacy in unselected DLBCL patients, pembrolizumab combined with R-CHOP was safe and associated with a high CR rate and improved 2-year PFS^58, 82^. BTLA has been reported to mark a high-checkpoint-expressing T-cell subset (PD-1, TIM-3, LIGHT, and LAG-3) with decreased cytolytic function and increased proliferation ability, thus correlating poor prognosis in DLBCL^13,83^.

## Conclusion

Taken together, the IPI-IPM risk score was compatible with the ability of tumor-infiltrating immune cells to determine the expression of immune checkpoints, suggesting that the poor prognosis of high-risk group may be due to the stronger immunosuppressive TME, and high-risk patients will benefit more from immune checkpoint inhibitors than low-risk patients, thereby resulting in a better prognosis. Our research provides new insights into the TME and immune-related therapies for DLBCL. However, it is noteworthy that some limitations came out because the conclusion was drawn from data from retrospective studies, and prospective studies are warranted to further confirm our results. In addition, functional and mechanistic studies of the genes in our risk model should be conducted to support their clinical application.

## Abbreviations

DLBCL: Diffuse large B-cell lymphoma
NHL: non-Hodgkin B-cell lymphoma
GCB Subtype: germinal center B-cell-like subtype
ABC Subtype: activated B-cell-like subtype
GEP: gene expression profiling
COO: cell-of-origin
SCNA: somatic copy number alterations
SV: structural variants
IPI: International Prognostic Index
TME: tumor microenvironment
ICB: immune checkpoint blockade therapy
GC form: germinal center-like form
MS form: mesenchymal form
IN form: inflammatory form
DP form: depleted form
WGCNA: weighted gene co-expression network analysis
IPI-IPM: IPI-based immune prognostic model
TCGA: the Cancer Genome Atlas
VST: variance stabilizing transformation
LME: DLBCL/Lymphoma microenvironment
DEGs: differentially expressed genes
GSEA: gene set enrichment analysis
GO: Gene Ontology
KEGG: Kyoto Encyclopedia of Genes and Genomes
NES: normalized enrichment score
OS: overall survival
Lasso: least absolute shrinkage and selection operator
AIC: Akaike information criterion
ROC: receiver operating characteristic
C-index: concordance index
DCA: decision curve analysis
t-SNE: t-distributed stochastic neighbor embedding
PCA: principal component analysis
GSVA: gene set variation analysis
ECM: extracellular matrix
TFs: transcription factors
Tregs: regulatory T cells
M0: non-activated macrophages
mDCs: myeloid dendritic cells
HSCs: hematopoietic stem cells
CAFs: cancer associated fibroblasts
F^GES^: functional gene expression signatures
MOA: mechanisms of action
FGF: fibroblast growth factor
Breg: regulatory B cells
IRAK: IL-1 receptor-associated kinase
ITAM: immunoreceptor tyrosine-based activation motif
PGE2: prostaglandin E2
PDTX: patient-derived tumor xenografts
DNMT: DNA methyltransferase
HAT: Histone acetyltransferase

## Declaration

### Ethical Approval and Consent to participate

Not applicable

### Consent for publication

All the authors read and approved the final version of the manuscript, as well as consented for publication.

### Availability of supporting data

The datasets used and/or analyzed during the current study are available from the corresponding author on reasonable request.

## Authors’ contributions

SM, DS, CS and YH conceived and designed the study. SM and DS did the literature research, performed study selection, data extraction, statistical analysis and wrote the draft. LA, FF and FP participated in the extraction and analysis of data. All the authors read and approved the final version of the manuscript.

## Acknowledgements

The results here are based upon data generated by the TCGA Research Network. The study reported herein fully satisfies the TCGA publication requirements (https://www.cancer.gov/tcga). The authors would like to thank the TCGA database developed by National Institutes of Health.

