## Supplementary figures and images for "An IPI based immune prognostic model for diffuse large B-cell lymphoma"

### Supplemental Figures

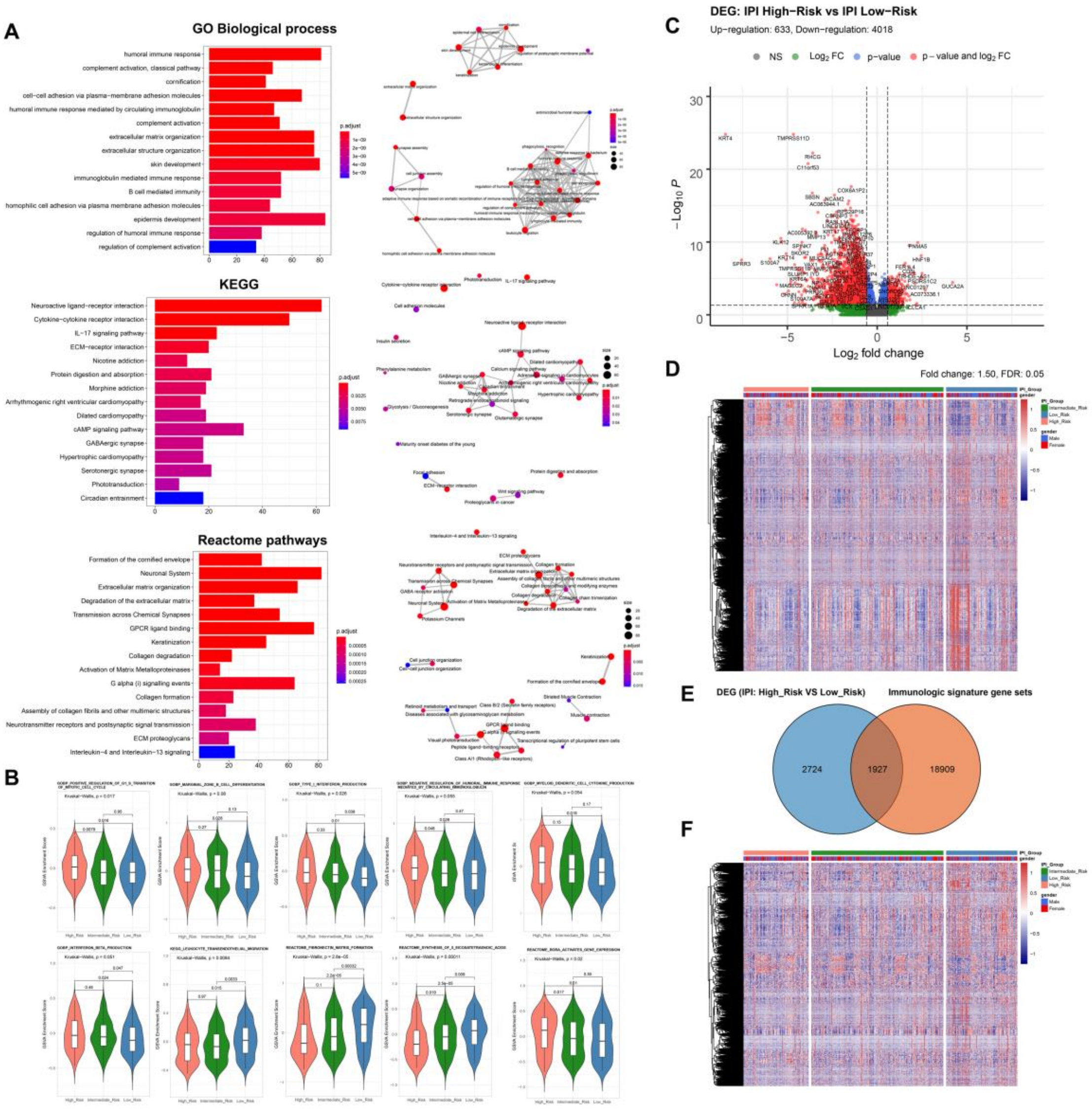

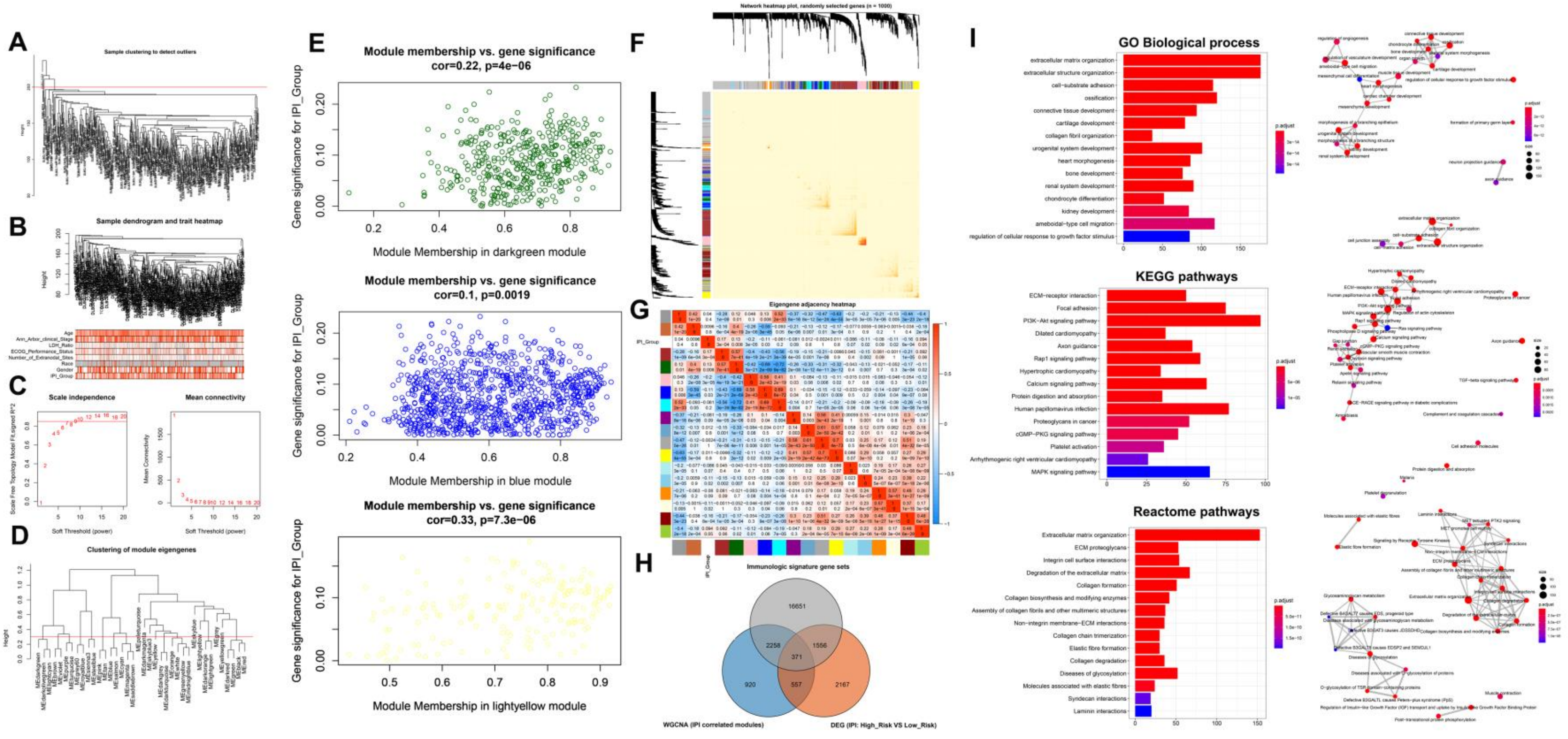

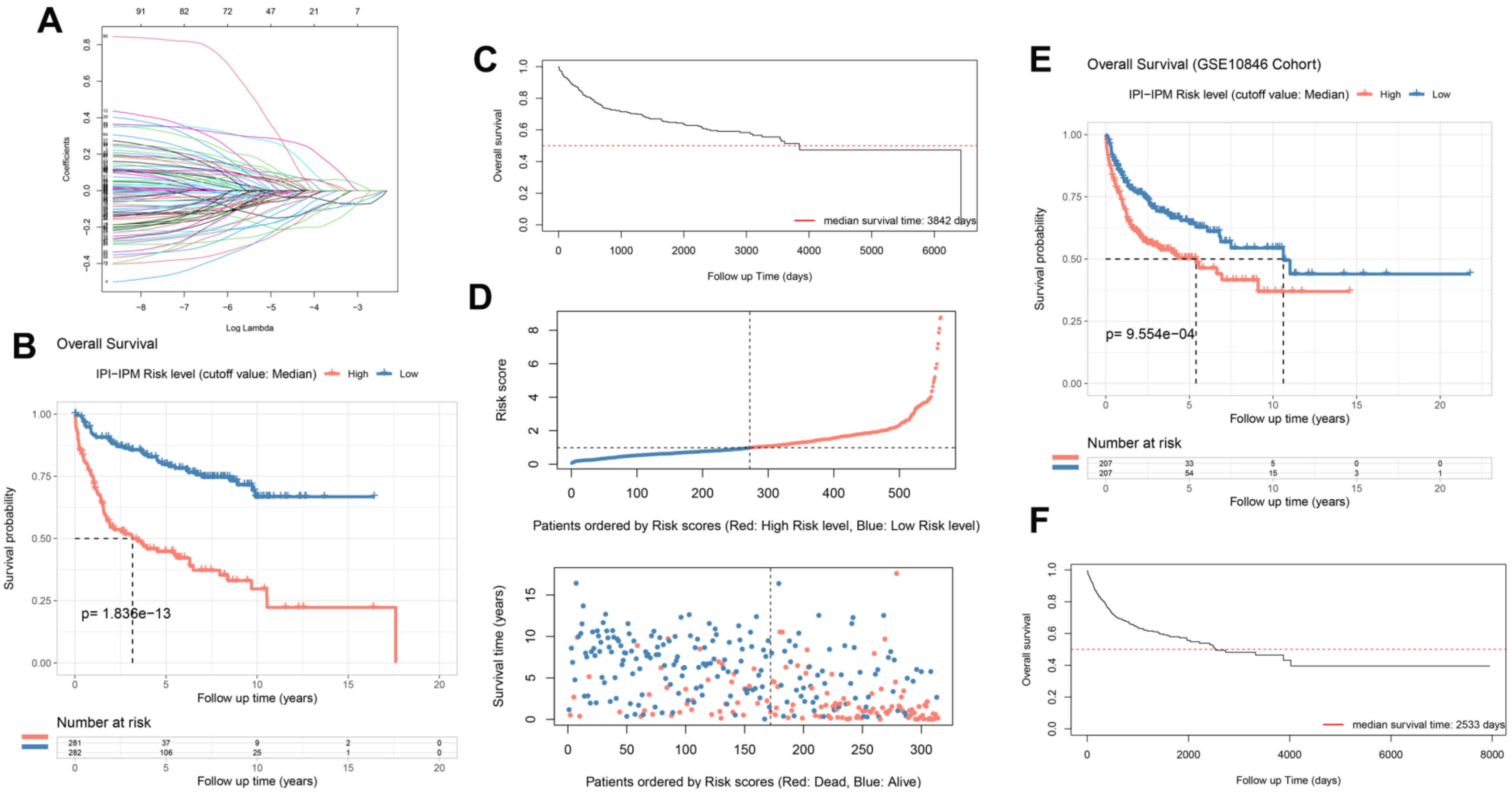

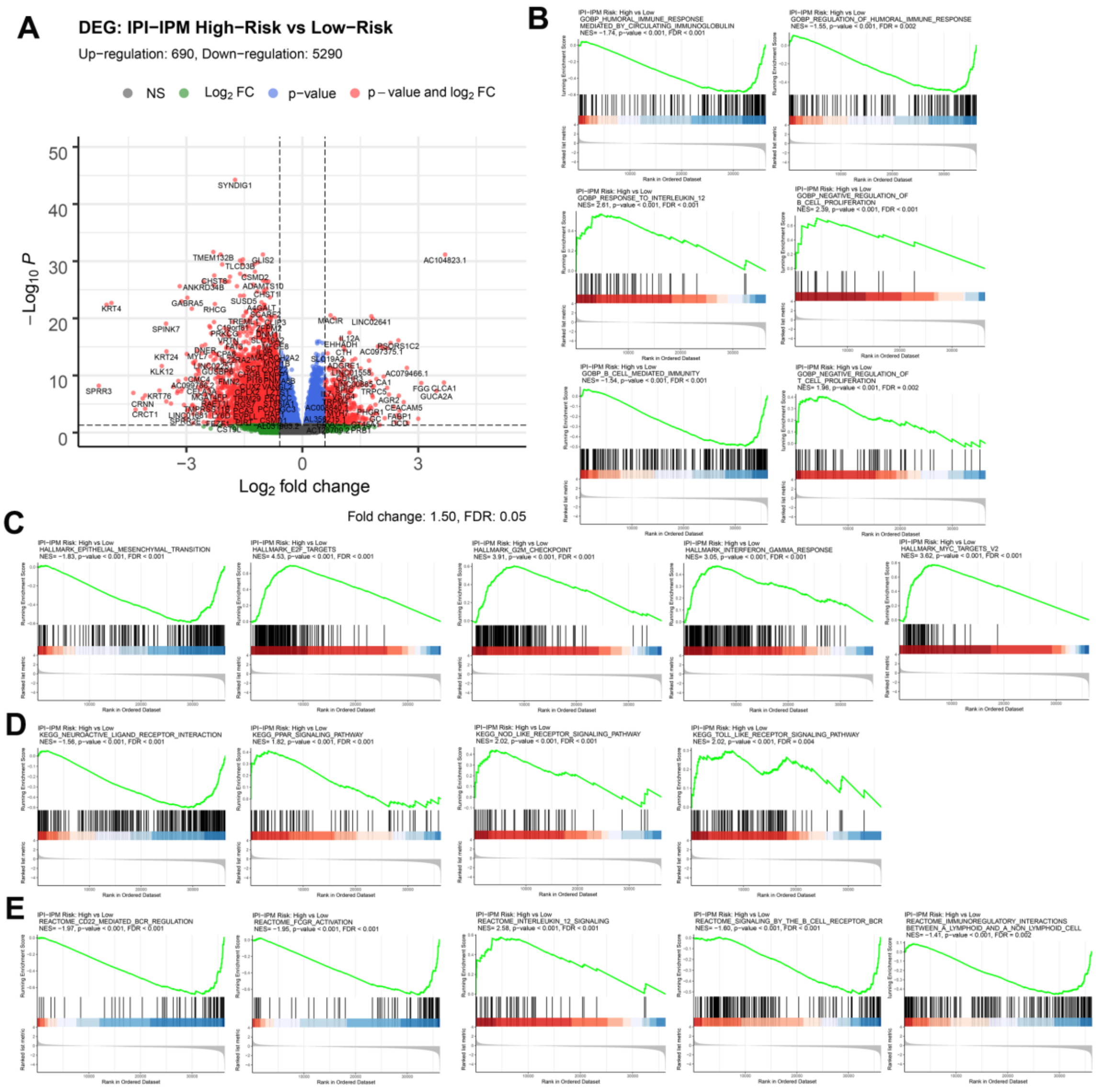

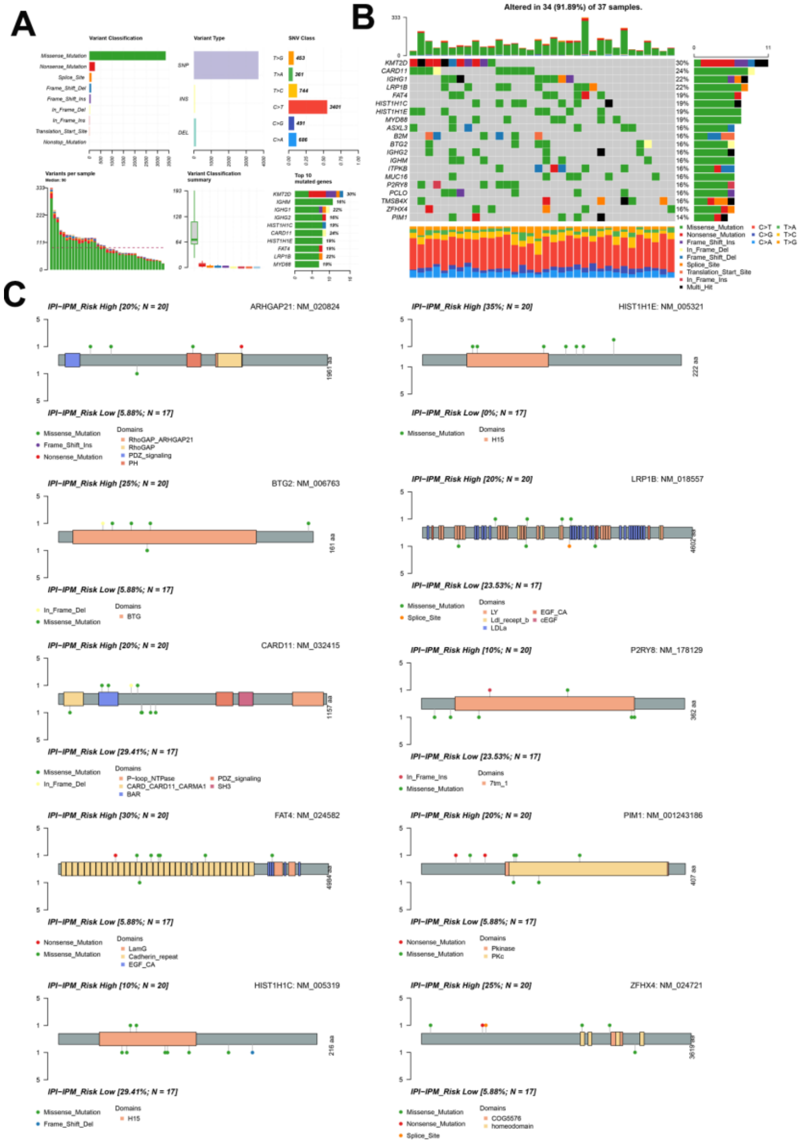

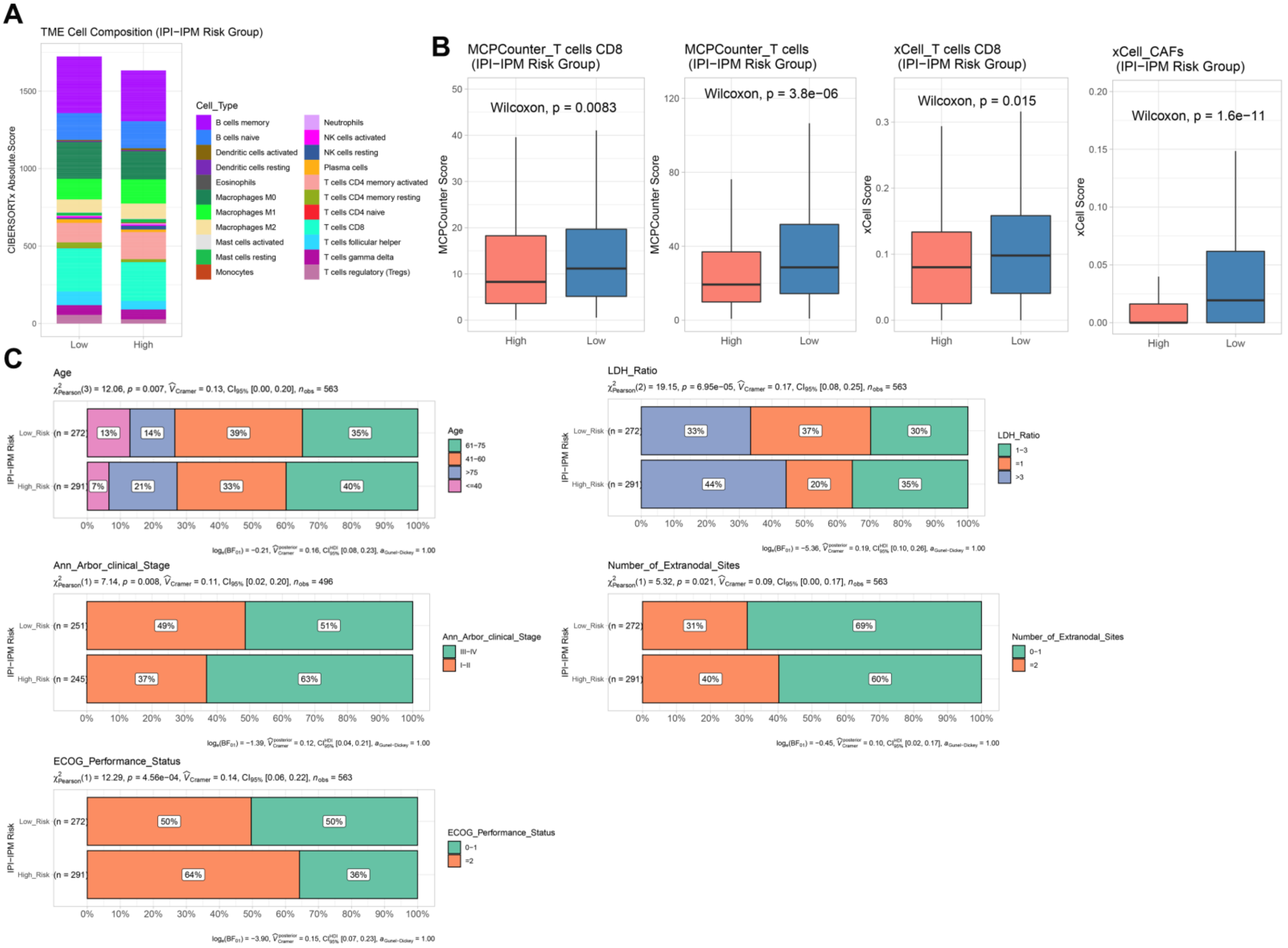
